# Frequency magnification in human primary auditory cortex nearly predicts behavioural frequency hyperacuity

**DOI:** 10.64898/2026.06.17.732895

**Authors:** Benjamin Gurer, Rosa-Maria Sanchez-Panchuelo, Susan Francis, Denis Schluppeck, Katrin Krumbholz, Julien Besle

## Abstract

The auditory system decomposes sounds into different frequencies, but does not represent them equally: in the human cochlea, lower frequencies occupy a larger spatial extent than higher frequencies, creating a quasi-logarithmic frequency map with finer low-frequency resolution. It remains unclear whether the primary auditory cortex preserves this quasi-logarithmic frequency mapping, or further magnifies behaviourally relevant frequencies, analogous to the foveal over-magnification in primary visual cortex that is thought to underpin visual hyperacuity phenomena such as Vernier acuity. Here, we used ultra-high field functional magnetic resonance imaging to quantify frequency magnification in the primary auditory cortex of 20 human listeners. We compared cortical frequency magnification to predictions based on published estimates of either cochlear frequency resolution or behavioural frequency discrimination performance, a form of perceptual frequency hyperacuity. Cortical frequency magnification was better predicted by frequency discrimination performance than by cochlear frequency resolution, with an additional unexpected over-representation of frequencies around 1 kHz. These findings suggest that frequency discrimination is constrained by cortical processes rather than by frequency information available at the cochlea, consistent with similar observations in the visual and tactile systems.

## Introduction

The auditory cortex contains frequency maps, in which nearby cortical locations preferentially respond to similar frequencies, and preferred frequency changes from low to high along a single spatial direction (Saenz and Langers, 2014; Schreiner and Winer, 2007). These cortical tonotopic (or cochleotopic) maps reflect the spatial organisation of the cochlea, which decomposes sounds into components of increasing frequencies from apex to base. While tonotopic organization is maintained from cochlea to cortex, little is known about whether, or to what degree, the internal structure of the map changes along the auditory processing hierarchy. Do cortical tonotopic maps faithfully preserve cochlear frequency mapping, devoting the same relative amount of cortical territory to different frequencies or do they over-represent – magnify – certain frequencies? Here we use ultra-high field (7 Tesla) functional magnetic resonance imaging (fMRI) to obtain detailed tonotopic maps in the human primary auditory cortex and quantify cortical frequency magnification functions.

Cochlear frequency mapping has been well characterised in both animal models and humans, showing characteristic frequency increases approximately exponentially with cochlear position from apex to base (Clopton et al., 1974; Greenwood, 1990). Since characteristic frequency changes more slowly with cochlear position at low than high frequencies, low frequencies occupy a larger proportion of the cochlea. If cortical maps preserve cochlear frequency mapping, they should exhibit a similar low-frequency magnification, such that the cortical frequency mapping function (relating cortical position along the tonotopic gradient to preferred frequency) is a scaled version of the cochlear frequency mapping function. In non-primate animal models, the cortical mapping function indeed broadly resembles that of the cochlea (Imaizumi and Schreiner, 2007; Kalatsky et al., 2005; Lee et al., 2004; Merzenich et al., 1973; Nishimura and Song, 2014; Robertson and Irvine, 1989; Storace et al., 2011), but the two mapping functions have not been quantitatively compared in any detail. In addition, the cortical frequency mapping function has not been measured in primates, including humans.

Evidence from other sensory modalities suggests that cortical and peripheral mapping functions can differ. Like the auditory system, both the visual and somatosensory systems are topographically organized, containing ordered maps of their respective perceptual spaces (retinal or skin location, respectively, analogous to frequency in hearing). As in the auditory system, these maps magnify specific regions of sensory space (Catani, 2017; Engel et al., 1997; Penfield and Boldrey, 1937; Sereno et al., 1995), reflecting systematic variations in peripheral receptor organisation – for example, increased photoreceptor and ganglion cell density in the fovea (Rolls and Cowey, 1970; Curcio and Allen, 1990; Wässle et al., 1990) and increased mechanoreceptor density on the skin of the hands and face (Darian-Smith and Kenins, 1980; Johansson and Vallbo, 1979). Crucially, in both visual and somatosensory systems, peripheral receptor density and cortical magnification functions do not align perfectly, and instead underpin different aspects of sensory performance: peripheral receptor density determines two-point acuity, that is, the ability to resolve two stimuli as separate (vision: Rolls and Cowey, 1970; touch: Brown, 2004; Craig and Lyle, 2001; Stevens and Choo, 1996; Van Boven and Johnson, 1994), whereas performance in other spatial judgement tasks may be primarily constrained by cortical magnification, rather than peripheral resolution. In both sensory modalities, positional discrimination performance (e.g. Vernier acuity, the judgement of alignment of two spatially separate stimuli) substantially exceeds two-point acuity, implying spatial precision far finer than predicted by peripheral receptor spacing (Geisler, 1984; Loomis, 1979; Westheimer, 1977). This hyperacuity is thought to result from population-level neuronal computations implemented centrally (Altes, 1989; Barlow, 1981; Duncan and Boynton, 2003; Levi et al., 1985; Westheimer, 2009), which would take place centrally. Indeed, variation in hyperacuity performance across sensory space is thought to more closely follow the cortical magnification function than peripheral receptor density (Duncan and Boynton, 2007, 2003; Levi et al., 1985; McKee et al., 1990; Westheimer, 1982). In the primary visual cortex, for instance, the fovea is more magnified than would be predicted by retinal cell density (Cowey and Rolls, 1974; Dow et al., 1981; Drasdo, 1977; Perry and Cowey, 1985; Van Essen et al., 1984; Azzopardi and Cowey, 1993, but see Curcio and Allen, 1990; Wässle et al., 1990). If the auditory system shares functional organisation principles with the visual and somatosensory systems (King and Nelken, 2009; Shamma, 2001), the cochlear and cortical frequency magnification functions may differ, and be associated with different types of frequency acuity.

While the terms acuity and hyperacuity are rarely used for hearing, there are clear parallels between frequency resolution (or selectivity) and two-point acuity on the one hand, and between frequency discrimination and hyperacuity on the other (Altes, 1989; Cariani and Micheyl, 2012). Frequency resolution is the degree to which frequency components are processed independently within the auditory system, and is commonly characterised as the inverse of auditory filter width – the frequency bandwidth over which masking occurs. Like visual and tactile two-point acuity, perceptual frequency resolution is constrained by peripheral properties, in this case the frequency selectivity of the cochlea, which, in turn, is determined by the relative spatial allocation, or magnification, of frequencies along the cochlea (Greenwood, 1990; Moore, 1986). In contrast, frequency discrimination performance, traditionally quantified by difference limens for frequency (DLFs), is not predicted by peripheral resolution limits: like visual and tactile hyperacuity, DLFs are much finer than auditory filter widths at the same frequency (by up to an order of magnitude or more; Moore, 1973; Wier et al., 1977), and also exhibit a different dependence on frequency: whilst auditory filter width increases approximately linearly with frequency, implying a corresponding decline in both frequency resolution and cochlear magnification, DLFs show an accelerating dependence on frequency (Fig. 1A). Consequently, frequency discrimination is hyperacute only at frequencies up to a few kHz, with DLFs increasing rapidly at higher frequencies to approach the width of auditory filters (Moore and Ernst, 2012; Nelson et al., 1983; Sek and Moore, 1995). A common interpretation is that hyperacute frequency discrimination at lower frequencies is not limited by place-based cues derived from the cochlear excitation pattern, but instead benefits from additional “temporal fine structure” cues conveyed by phase-locking of auditory nerve response (Moore, 1973; Moore and Glasberg, 1989; Sek and Moore, 1995). Under this interpretation, the steep increase in DLFs would reflect the breakdown of phase-locking at high frequencies (Joris and Verschooten, 2013; Verschooten et al., 2019, 2018). An alternative explanation, however, is that frequency discrimination, as a form of hyperacuity, is not primarily constrained by cochlear processes, but instead reflects population-based computations in more central auditory structures such as the primary auditory cortex, in analogy to visual or tactile hyperacuity (Altes, 1989; Cariani and Micheyl, 2012; Micheyl et al., 2013; Verschooten et al., 2019). If this interpretation is correct, frequency magnification in primary cortical tonotopic maps should reflect the frequency dependence of DLFs, rather than the cochlear frequency resolution function.

**Figure 1:**
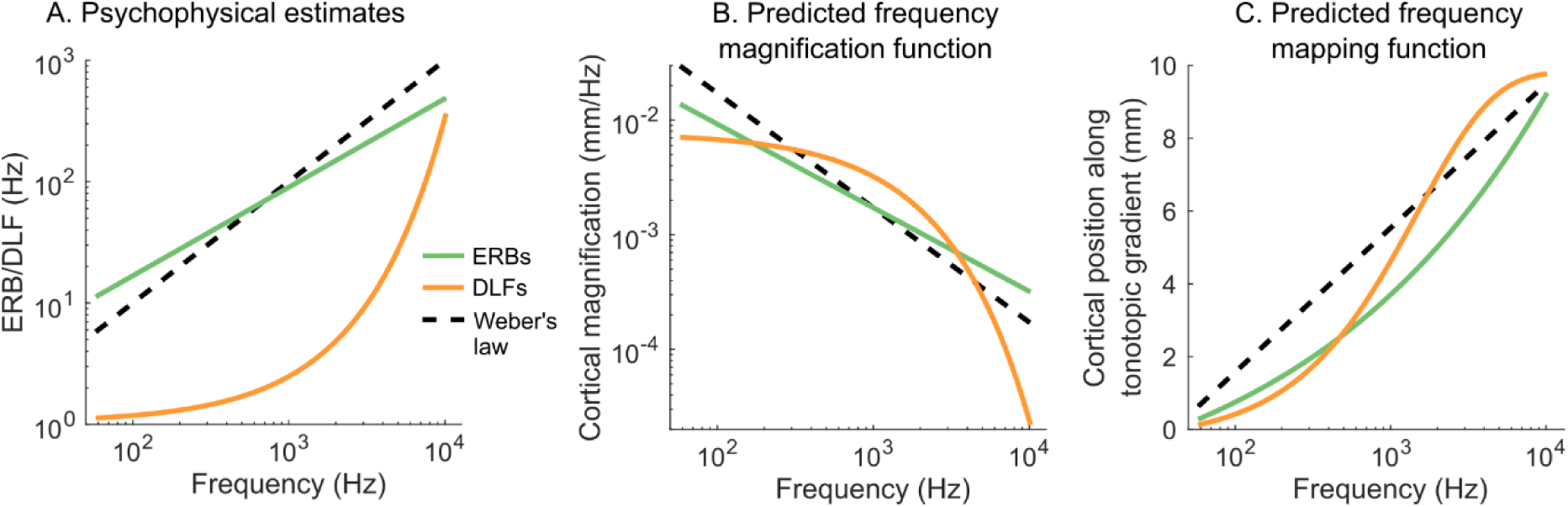
Psychophysical estimates of cochlear filter width and frequency discrimination thresholds and corresponding predictions for cortical frequency magnification and mapping functions. A. Idealised functions relating cochlear filter widths, measured through notched-noise forward masking and expressed as equivalent rectangular bandwidths (ERBs; green line; Oxenham and Shera, 2003), and pure-tone difference limens for frequency (DLFs; orange line, Micheyl et al., 2012), to stimulus frequency. For comparison, the dashed black line depicts a hypothetical case in which filter width is a fixed Weber fraction (10%) of frequency. Note the logarithmic scaling of both axes. B. Reciprocals of the ERB and DLF functions from A, representing the so-called ERB- and DLF-rate functions (the number of consecutive ERBs or DLFs per Hz) which are proportional to cortical frequency magnification (in mm/Hz), up to an unknown scaling factor. Each function was multiplied by a constant factor (in mm/ERB or mm/DLF) to allow comparison with fMRI-based measurements (see Fig. 3E); only relative shapes, not absolute value, can be meaningfully compared between predictions. C. Predicted frequency mapping functions, obtained by integrating the magnification functions in B with respect to frequency. Because frequency is plotted in logarithmic units, a steeper slope indicates greater magnification relative to Weber’s law.

Here we measure the frequency magnification functions of two tonotopic maps in the human primary auditory cortex and compare their shape to predictions based on the known frequency dependence of either cochlear frequency resolution or perceptual frequency discrimination performance, quantified as DLFs. We find that frequency magnification in both cortical maps is better predicted by DLFs than by cochlear frequency resolution, suggesting both that cortical and cochlear frequency magnification functions differ and that frequency discrimination is a form of perceptual hyperacuity analogous to Vernier acuity in vision, constrained by central, perhaps cortical, population-level processes. In addition, cortical frequency magnification shows an unexpected over-representation of frequencies around 1 kHz compared to predictions based on the frequency discrimination performance.

## Results

### Cortical frequency mapping is better predicted by frequency hyperacuity than frequency resolution

To test whether cortical frequency mapping reflects cochlear organisation, or instead tracks frequency discrimination performance, we predicted frequency mapping and magnification functions from published estimates of either perceptual frequency resolution (taken to reflect cochlear frequency resolution and mapping) or frequency discrimination performance. Figure 1A compares estimates of frequency resolution, measured as the equivalent rectangular bandwidth (ERB) of auditory filters (green solid line) and frequency discrimination thresholds (DLFs; yellow solid line), both derived from psychophysical measurements in humans (Micheyl et al., 2012; Oxenham and Shera, 2003). ERBs and DLFs increase with frequency at different rates: cochlear ERBs increase slightly less than proportionally to frequency (compared to a strictly proportional relationship, or “Weber’s law”, dashed black line), while DLFs are substantially narrower than ERBs at low frequencies and increase progressively more steeply with frequency.

Hypothetical frequency magnification and mapping functions can be derived by assuming that measures of frequency resolution or discrimination (ERBs or DLFs), evaluated at any given frequency, correspond to a constant spatial distance along the tonotopic gradient (Greenwood, 1990; Moore, 1986). Under this assumption, predicted frequency magnification (the amount of cortical space dedicated to a particular frequency; Fig. 1B) is proportional to the reciprocal of the frequency-dependent resolution or discrimination shown in Fig. 1A, up to a scaling factor (expressed in ERB/mm or DLF/mm). The corresponding frequency mapping function, which maps the position along the tonotopic gradient to preferred frequency, is then obtained by integrating the magnification function with respect to frequency (Fig. 1C). Because ERBs and DLFs increase with frequency at different rates, the two models make distinct predictions for how cortical magnification declines with frequency (Fig. 1B), and consequently for the shape of the resulting frequency mapping functions (Fig. 1C). Under the ERB model, magnification decreases gradually with frequency, reflecting the slightly sub-proportional increase of ERBs relative to Weber’s law, and predicts a progressively steeper frequency mapping function. Under the DLF model, in contrast, magnification decreases more slowly at low frequencies than at high frequencies, predicting maximal relative magnification (compared to Weber’s law), and the steepest mapping between 1 and 2 kHz.

We evaluated these predictions against empirical fMRI measurements of frequency mapping in two tonotopic maps of the primary auditory cortex of 20 participants. Tonotopic (preferred frequency) maps were derived from blood-oxygen-level-dependent (BOLD) responses to narrowband noises at different centre frequencies (Fig. 2). In the first 12 participants, noises were presented at seven centre frequencies between 0.25 and 6 kHz, whereas, in the remaining eight participants, a denser set of 32 frequencies between 0.1 and 8 kHz was used (Fig. 2A). From the tonotopic maps, reconstructed on individual cortical surfaces (Fig. 2B), we identified two mirror-reversed tonotopic gradients in each hemisphere, hereafter referred to as the “anterior and posterior gradient regions of interest (ROIs)”. These two gradients were separated by a low-frequency gradient reversal located along the posterior half of Heschl’s gyrus, and were bounded anteriorly and posteriorly by high-frequency reversals. In all individual hemispheres, both gradients overlapped a region of increased frequency selectivity (Fig. 2B, right), likely corresponding to the primary auditory cortex (Besle et al., 2019; Da Costa et al., 2011).

**Figure 2:**
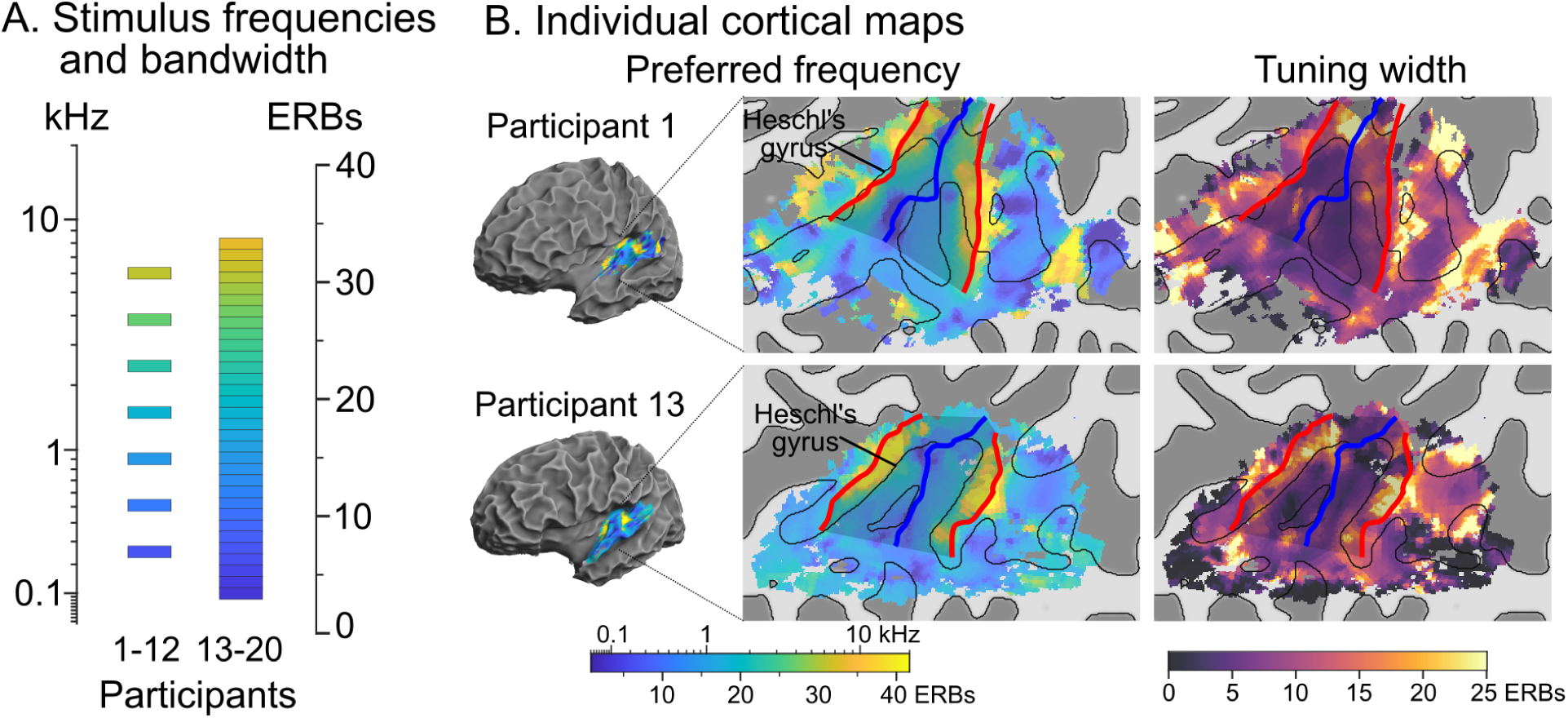
Auditory stimuli and cortical maps of preferred frequency and population tuning width. A. Narrowband noise (NBN) stimuli were presented at either 7 centre frequencies between 0.25 and 6 kHz (participants 1-12) or 32 centre frequencies between 0.1 and 8 kHz (participants 13-20). In both cases, centre frequencies were equally spaced on the ERB-number scale (Glasberg and Moore, 1990) and NBNs were one ERB wide. B. Voxelwise maps of preferred frequency (left) and tuning width (right), displayed on the reconstructed left-hemisphere cortical surface and flattened supratemporal plane of two participants (see Figure 2—figure supplements 1-2 for all participants and hemispheres). The preferred-frequency colour scale matches that used in A. Dark and light grey shading represent negative and positive curvature, respectively, and thin black lines curvature inflections (gyrus/sulcus boundaries). In all participants, two mirror-reversed tonotopic gradients were observed, separated by a low-frequency gradient reversal (thick blue line) and bounded anteriorly and posteriorly by high-frequency reversals (thick red lines). Reversals, identified using an automated, unbiased procedure (Schönwiesner et al., 2015) were used to define the anterior and posterior gradient ROIs (darker shading between the reversals). Tuning width maps (rightmost panel) show that both gradient ROIs overlapped with a region of high frequency selectivity (low tuning width), consistent with primary auditory cortex (Besle et al., 2019; Da Costa et al., 2011).

Within each gradient ROI (80 in total), we characterized frequency mapping using three alternative measurements, which we then compared to the ERB and DLF predictions shown in Figure 1. First, we measured the magnitude of the local preferred frequency gradient vector at each point of the cortical surface (Fig. 3A), reflecting the rate at which preferred frequency changes with cortical position. Gradient magnitude (in Hz/mm) is the reciprocal of frequency magnification (in mm/Hz); accordingly, the function relating gradient magnitude to preferred frequency (Fig. 3D) can directly be compared to the ERB and DLF functions shown in Fig. 1A, up to a scaling factor in ERB/mm or DLF/mm. Second, we measured the surface area within the ROI preferring a given frequency (Fig. 3B), which is a direct estimate of areal frequency magnification (the product of linear magnification along the tonotopic gradient by the average length of the iso-frequency contour within the gradient region). When plotted as a function of preferred frequency (Fig. 3E), this measure can be compared to the ERB- and DLF-based magnification predictions shown in Fig. 1B, up to a scaling factor in mm^2^/ERB or mm^2^/DLF. Third, we quantified frequency mapping by estimating the position of each surface vertex along the tonotopic progression within a given ROI, defined as the vertex’s shortest path length along the cortical surface to the low-frequency gradient reversal separating the two gradient ROIs (Fig. 3C). Plotting cortical distance as a function of vertex preferred frequency (Fig. 3F) yields an empirical frequency mapping function that can be compared to the ERB and DLF-based predictions shown in Fig. 1C, again up to a scaling factor in ERB/mm or DLF/mm.

**Figure 3:**
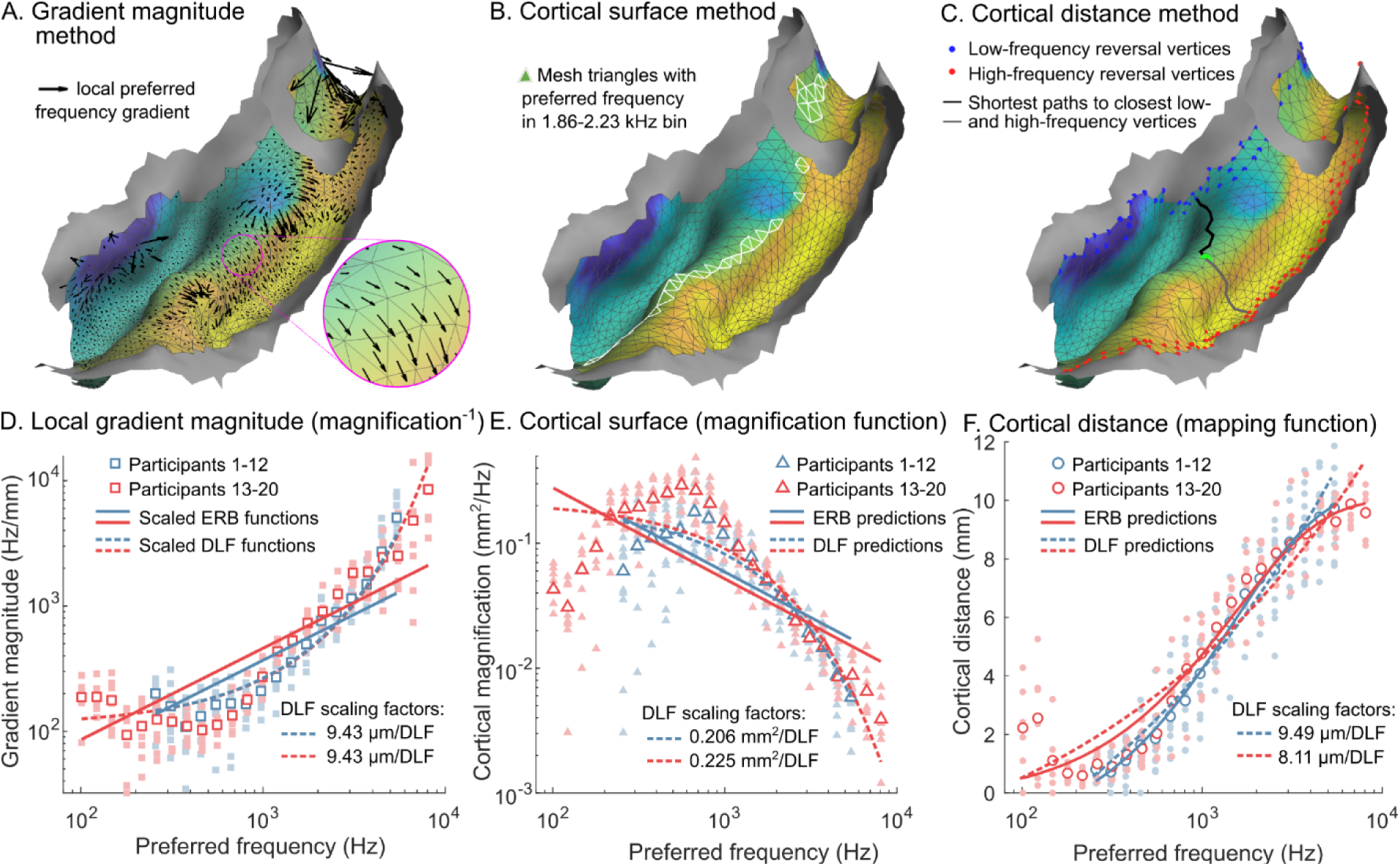
Three fMRI-based measures used for testing the ERB- and DLF-based predictions of frequency magnification and tonotopic mapping. (A-C) Illustration of how the three measures were derived, for the left posterior gradient ROI of participant 1. (D-F) Corresponding group results plotted as a function of preferred frequency and compared to best-fitting predictions. A. Local preferred frequency gradients were computed at the center of each surface mesh triangle within each individual gradient ROI. Arrow length indicates gradient magnitude in octaves/mm. B. Cortical surface area of cortex representing a given frequency was quantified as the summed surface area of mesh triangles whose preferred frequency fell within one of 24 log-spaced preferred frequency bins between 0.1 and 8 kHz (highlighted triangles fall in the 1.86-2.23 kHz bin). C. Cortical position along the ROI’s tonotopic axis was computed as the shortest Dijkstra distance between each vertex (example shown in bright green) to the closest vertex of the low-frequency gradient reversal (blue dots) and normalized by total distance between the closest low- and high-frequency reversal vertices (red dots). D. Median local gradient magnitude in each preferred frequency bin, averaged across both hemispheres and both anterior and posterior gradient ROIs and plotted as a function of the bins’ centre frequency. Results from participants 1-12 and 13-20 are shown in blue and red, respectively. Open symbols indicate group averages and filled symbols individual data. Solid and dashed lines depict the best-fitting (scaled) ERB- and DLF-based predictions for each participant group. The scaling factors for the DLF fits are reported at the bottom of the panel. E. Cortical surface area as a function of each bin’s centre frequency, plotted using the same conventions as in D. F. Median cortical distance along the tonotopic gradient in each bin, as a function of the bins’ centre frequency, again plotted using the same conventions as in D.

Figure 3D-F shows the three fMRI-derived empirical functions – local frequency gradient magnitude, areal frequency magnification and shortest cortical distance), each plotted as a function of preferred frequency and averaged across hemispheres and gradient ROIs – and compares them to the best-fitting (scaled) ERB- and DLF-based predictions. The DLF-based predictions provided a substantially better fit than the ERB predictions across all three measures, and in both groups of participants. The magnitude of the local frequency gradient (Fig. 3D) remained approximately constant (or slightly decreased) at low frequencies up to ∼0.6 kHz, before increasing more than linearly with frequency, closely resembling the DLF, rather than ERB, frequency function. Bayesian model comparison using the Bayesian information criterion approximation provided very strong evidence in favour of the DLF model in both groups (participants 1-12: BF_DLF,ERB_ = 1.81 × 10^8^; participants 13-20: BF_DLF,ERB_ = 3.75 × 10^4^). Empirical magnification functions derived from cortical surface area (Fig. 3E) increased with frequency up to ∼0.6 kHz, and then decreased more than inversely with frequency, again more closely matching the DLF-based prediction. Bayes factors strongly favoured the DLF model in both groups (participants 1-12: BF_DLF,ERB_ = 5909; participants 13-20: BF_DLF,ERB_ = 781). Finally, the empirical frequency mapping function obtained from cortical distance (Fig. 3F), exhibited a sigmoidal shape with its steepest slope around 1 kHz, consistent with the DLF-predicted mapping function. Bayesian model comparison again strongly favoured the DLF prediction in both groups (participants 1-12: BF_DLF,ERB_ = 21.8; participants 13-20: BF_DLF,ERB_ = 505). Similar results were obtained when the measurements were averaged separately in each hemisphere or in each gradient ROI (see Fig. 3—figure supplement 1).

The scaling factors relating the DLF predictions to the empirical cortical functions are reported at the bottom of Figure 3 for each measurement and participant group. For the gradient magnitude and cortical distance measures, scaling factors are expressed in mm/DLF and correspond to the average cortical distance along the tonotopic gradient associated with a one-DLF change in preferred frequency. In contrast, scaling factors derived from cortical surface area are expressed in mm^2^/DLF and represent the cortical area devoted to a one-DLF frequency range. These areal scaling factors reflect the linear scaling factor multiplied by the average length of iso-frequency contours orthogonal to the tonotopic gradient. Areal scaling factors were slightly greater than 0.2 mm^2^/DLF, while linear scaling factors were around 8-9 μm/DLF. Together, these values imply iso-frequency contour length of 23 mm on average, approximately corresponding to the extent of the gradient ROIs along Heschl’s gyrus.

### Cortical frequency magnification around 1 kHz is larger than predicted by frequency hyperacuity performance

Although cortical frequency mapping functions were better predicted by frequency discrimination performance (DLFs) than frequency resolution (ERBs), the DLF model did not fully capture their shape (see Fig. 3D-F). To evaluate departures from the DLF prediction, we derived the frequency magnification function from each of the three empirical measures (Fig. 4A-C). Frequency magnification was derived from local gradient magnitude by taking its reciprocal (Fig. 4A) and from the cortical-distance mapping function by differentiating with respect to preferred frequency (Fig. 4C). Across all three measures, the resulting magnification functions showed similar systematic deviations from the DLF prediction: frequency magnification was smaller than predicted by the best-fitting DLF model at low frequencies (< 0.3 kHz), but larger than predicted at frequencies just below or around 1 kHz (depending on the measure). We statistically evaluated these departures by re-expressing frequency magnification in units of mm/DLF (Fig. 4D-F), obtained by multiplying magnification (in mm/Hz) by the corresponding DLF size (in Hz) at each frequency (see Methods). If cortical frequency magnification followed the DLF prediction exactly, it would be inversely proportional to DLF size, and therefore constant when expressed in mm/DLF, at a value given by the best-fitting DLF scaling factor (reported at the bottom of Fig. 3D-F). Instead, cortical magnification in mm/DLF varied markedly with frequency, showing a consistent pattern across all three measures, with values varying by nearly an order of magnitude from a minimum at low frequencies and a pronounced maximum just below or around 1 kHz.

**Figure 4:**
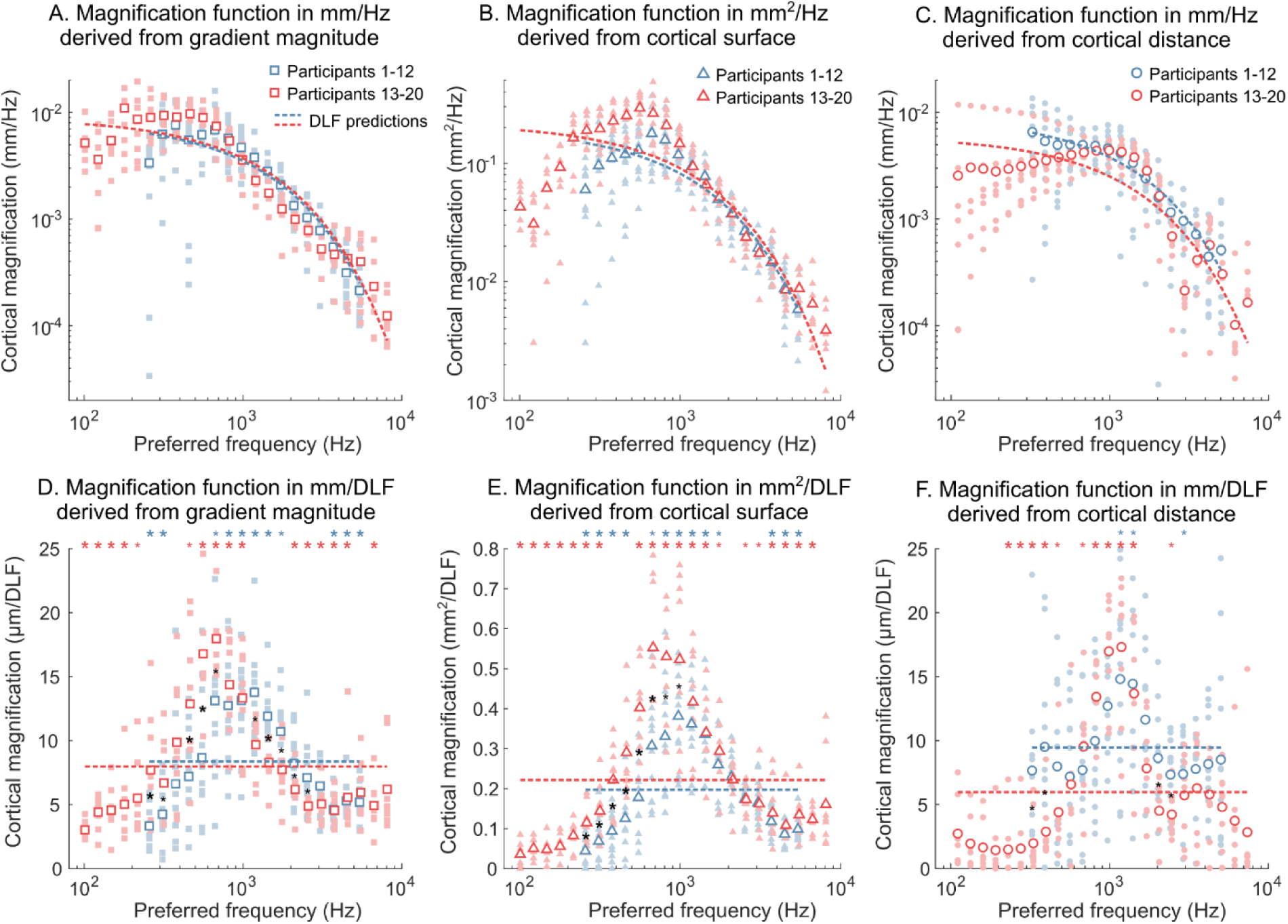
Frequency magnification functions in primary auditory cortex, expressed in linear (Hz) or DLF-based frequency units. A-C) Frequency magnification functions derived from local preferred frequency gradient magnitude (A), cortical surface area (B) or cortical distance measurements (C). Plotting conventions are the same as in Fig. 3D-F. Dashed lines show the best-fitting DLF-based predictions for each participant group (blue: participants 1-12; red: participants 13-20). (D-F) Same as A-C, but magnification expressed in mm/DLF (or mm^2^/DLF for the surface-derived function; E) rather than mm/Hz (or mm^2^/Hz). Small and large coloured star symbols mark preferred frequency bins with at least moderate evidence (Bayesian t-tests, BF_10_ > 3) that magnification differed from the best-fitting DLF prediction, before and after correction for multiple comparisons across bins, respectively. Small and large black star symbols indicate bins with at least moderate evidence for group differences.

To formally test these departures from the DLF prediction, we fitted three mixed-effects Bayesian ANOVA models – one for each measure – to the frequency magnification values expressed in mm^(2)^/DLF. Each model included preferred frequency, group (participants 1-12 vs 13-20), gradient ROI (anterior vs posterior) and hemisphere (left vs right) as fixed factors, and participant as a random factor. All three analyses provided very strong support for a main effect of preferred frequency (gradient magnitude: BF_10_ = 1.52 × 10^100^; cortical surface: BF_10_ = 1.84 × 10^194^; cortical distance: BF_10_ = 1.66 × 10^33^), confirming that cortical frequency magnification in mm/DLF did not remain constant across frequency. In addition, all three analyses provided very strong evidence for an interaction between preferred frequency and group (gradient magnitude: BF_10_ = 1.48 × 10^25^; cortical surface: BF_10_ = 6.25 × 10^18^; cortical distance: BF_10_ = 2.50× 10^5^). This interaction reflects a modest shift in the frequency at which cortical magnification reached its maximum, with the peak occurring around 1 kHz in participants 1-12 but slightly below 1 kHz in participants 13-20. Despite this shift, the overall pattern of departures was highly similar across groups for all three measures. Post-hoc Bayesian T tests at each frequency bin provided moderate to very strong evidence that cortical magnification exceeded the DLF-based prediction at several frequency bins around or just below 1 kHz, but that the opposite was true at both lower and higher frequencies, after correction for multiple comparisons across frequency bins (see coloured star symbols in Fig. 4D-F).

The same Bayesian ANOVAs also provided evidence for modest differences in frequency magnification across gradient ROIs and hemispheres, but these effects were not consistently observed across all three measures. Specifically, both the gradient magnitude and cortical surface measures provided evidence that magnification was, on average, 10-15% larger in the right than the left hemisphere. Cortical surface measures also provided evidence that magnification was up to 90% larger in the anterior than the posterior gradient ROI at a limited number of frequency bins above 2kHz. Gradient magnitude showed a similar effect in participants 1-12 only, which did not survive correction for multiple tests. In contrast, the cortical distance measure provided evidence against differences between both hemispheres or gradients. Full statistical results are provided in Figure 4—figure supplement 1.

## Discussion

Here we show that the frequency mapping of human primary auditory cortex is better explained by behavioural frequency discrimination limits than by the cochlear frequency map and associated frequency resolution limits. We found that cortical magnification varies systematically with frequency discrimination performance, such that magnification is approximately inversely proportional to behavioural difference limens (DLFs), mirroring the relationship between cortical magnification and perceptual acuity in the visual system. This relationship is not, however, exact. While behavioural thresholds explained a substantial proportion of the variance in cortical magnification, we observed a consistent over-representation of frequencies around 1 kHz relative to this prediction. The origin of this deviation is currently unclear, but may reflect additional functional specialisation, the particular ecological relevance of this frequency range in speech and music, and/or residual methodological or modelling factors not captured by the current framework. Importantly, our findings were robust across three independent approaches to estimating cortical magnification functions, underscoring the reliability of the observed relationship between cortical representation and perceptual discrimination of sound frequencies.

### Frequency mapping and magnification in primary auditory auditory cortex compared to the cochlea

In the cochlea, frequency mapping is approximately logarithmic: characteristic frequency increases slightly less than exponentially from apex to base, and, consequently, both frequency resolution and magnification decrease progressively (slightly less than linearly) from low to high frequency (Clopton et al., 1974; Greenwood, 1990; Moore, 1986). In contrast, we show – using three alternative estimation methods – that cortical magnification remains approximately constant at low frequencies (or perhaps even increases slightly with frequency), before starting to decrease steeply with frequency between 0.6 and 1kHz (depending on the measurement method). This divergence between cochlear and cortical magnification functions suggests a relative over-representation of mid-range frequencies in cortex, most prominently around 0.6–1 kHz, near the inflection point of the cortical magnification function, implying a redistribution of representational resources along the auditory pathway.

This observation contrasts with previous invasive measurements in non-primate mammalian auditory cortex, which have generally suggested a quasi-logarithmic cortical frequency map resembling that of the cochlea (Imaizumi and Schreiner, 2007; Mrsic-Flogel et al., 2006; Nishimura and Song, 2014; Robertson and Irvine, 1989). While some animal studies have reported deviations between cochlear and cortical mapping, these typically involved an over-representation of high frequencies rather than mid-frequencies as observed in the current study (Imaizumi and Schreiner, 2007; Merzenich et al., 1975; Nishimura and Song, 2014). This suggests that discrepancies between cochlear and cortical maps may reflect species-specific ecological or communicative demands, reflecting a redistribution of cortical resources towards behaviourally relevant parts of the spectrum. Given that prior animal work has employed a wide range of recording techniques (including electrophysiology and optical imaging) and stimulus levels (both near- and supra-threshold), it seems unlikely that the observed interspecies differences can be fully explained by methodological factors. Nevertheless, direct comparisons using fMRI tonotopic mapping in animal models, particularly in primates and under stimulation conditions comparable to those used here, will be important to resolve this discrepancy.

### Relationship between cortical magnification and frequency hyperacuity

Across sensory modalities, cortical magnification has been shown to reflect perceptual acuity rather than the physical distribution of receptors, as exemplified by the relationship between visual cortical magnification and Vernier acuity, and between somatosensory magnification and tactile discrimination (Azzopardi and Cowey, 1993; Duncan and Boynton, 2007; see introduction for further references). Our findings extend this principle to audition, showing that cortical frequency magnification follows behavioural frequency discrimination thresholds. This correspondence between cortical magnification and perceptual acuity has important potential implications for how frequency discrimination is implemented in the auditory system and for the type of information that constrains frequency discrimination performance. In this framework, cortical processing itself may constitute the limiting stage for auditory frequency discrimination.

This interpretation contrasts with the classical view that hyperacute frequency discrimination at low and mid-frequencies relies on the availability of temporal fine-structure cues in addition to rate-place cues (Moore, 1973; Moore and Glasberg, 1989; Sek and Moore, 1995). As noted previously (Micheyl et al., 2013; Verschooten et al., 2019), this traditional account assumes that rate-place cues are is read out locally, from a single cochlear channel, implying that purely place-based thresholds should not be smaller than the just-noticeable difference in intensity at a given frequency (approximately 1 dB, Zwicker, 1970). However, models that allow pooling of rate-place information across multiple cochlear locations show that the place cues alone can, in principle, support hyperacute frequency discrimination (Micheyl et al., 2013; Siebert, 1970), obviating the need for a temporal code (Lau et al., 2017; Verschooten et al., 2019, 2018). Such pooling is likely to occur at the cortical level (Micheyl et al., 2013), analogous to proposals in vision that hyperacuity arises from integration across multiple locations in primary visual cortex (Barlow, 1981; Duncan and Boynton, 2003; Levi et al., 1985; Westheimer, 2009). From this perspective, the shape of the frequency discrimination function may reflect constraints imposed by cortical organization – such as the frequency magnification function measured here – rather than to a change in the nature of the peripheral frequency code. As has been pointed out previously, cortical frequency magnification may be shaped by the ecological relevance of different frequencies (Lau et al., 2017; Verschooten et al., 2019). A clear example of such behaviourally driven over-representation is found in echolocating bats, in which a restricted range of echo-relevant frequencies is selectively magnified in the auditory pathway (Macias et al., 2020; Pollak and Casseday, 2012; Radtke-Schuller and Schuller, 1995; Schuller and Pollak, 1979). A notable difference is that, in bats, this “acoustic fovea” already exists in the cochlea (Pollak and Casseday, 2012; Vater et al., 1985), whereas the present results suggest that cortical over-magnification of frequencies around 0.6-1 kHz in humans emerges centrally and is not inherited from the periphery. It is unclear what specific aspect of behavioural relevance or environmental sounds could contribute to the observed cortical over-representation. Speech would seem an obvious candidate, but the most important frequencies for speech intelligibility, as measured by frequency importance function, tend to peak at ∼2kHz (DePaolis et al., 1996; Kates, 2013), rather than near 0.6-1 kHz. More generally, the presently observed pattern of over-magnification highlights a key difference from other sensory systems. In vision and touch, cortical magnification typically reinforces peripheral specialisations, with over-represented regions in cortex corresponding to areas already magnified at the receptor level (e.g., the fovea or fingertips; Azzopardi and Cowey, 1993; Duncan and Boynton, 2007). In contrast, auditory cortical magnification, as observed here, emphasises a frequency range that is not preferentially represented in the cochlea. This dissociation further supports the idea that cortical magnification is shaped less by peripheral encoding constraints and more by central processing demands, even when the specific ecological drivers are not immediately apparent. In the auditory domain, we cannot exclude the possibility that the expanded cortical representation of mid-frequencies reflects the availability of peripheral temporal fine-structure cues, which may be re-coded and integrated with place-based information within a unified cortical representation (Micheyl et al., 2013).

The involvement of auditory cortex in frequency discrimination is consistent with observations that both human patients and primate models with bilateral cortical lesions show substantial, and in some cases task-specific, increases in frequency discrimination thresholds (Dykstra et al., 2012; Harrington et al., 2001; Tramo et al., 2002). Moreover, frequency discrimination training in monkeys selectively expands the cortical representation of the trained frequencies (Recanzone et al., 1993). In contrast, non-primate mammals – who do not appear to exhibit a comparable over-representation of mid-frequencies in auditory cortex (see above) – show relatively limited deficits following bilateral cortical ablation (e.g., Gimenez et al., 2015, although this has not been tested near discrimination threshold, Tramo et al., 2002). Consistent with this, improvements in frequency discrimination can occur without clear tonotopic reorganisation in some species (e.g., cats: Brown, 2004), although such reorganisation has been observed in others (rats: Polley et al., 2006), or may have been transient (Reed et al., 2011). Taken together, these findings support the idea that auditory cortex plays a more central role in frequency discrimination tasks in humans – and possibly primates more generally – than in other mammals.

Although cortical magnification was better predicted by frequency discrimination performance than by cochlear magnification, discrimination performance did not fully account for the observed cortical mapping: the most strongly magnified frequencies were centred around or just below 1 kHz, rather than between 1 and 2 kHz as predicted by published estimates of frequency discrimination performance (see Fig. 1B). One possibility is that this mismatch stems from individual differences between the participant samples contributing to the cortical and behavioural measurements (see next section). Alternatively, it may indicate that the population code underlying frequency discrimination is not determined solely by the primary auditory cortex, but instead reflects contributions from other central auditory populations with slightly different magnification profiles. Given the multiple subcortical stages in the auditory pathway, cortical magnification could partially inherit its structure from earlier nuclei, such as the inferior colliculus. However, precise measurements of frequency mapping in subcortical auditory structures are scarce, both in humans and in animal models (with the notable exception of echolocating bats, e.g. Schuller and Pollak, 1979).

### Experimental factors affecting the measurements

It is important to keep in mind some of the limitations of the results reported here.

First, cortical magnification was compared to published estimates of frequency discrimination and cochlear tuning rather than measurements obtained in the same participants. While these published datasets are based on larger samples and are therefore likely more robust, individual variability could introduce a mismatch between behavioural and cortical measurements. This may contribute to the discrepancy between the observed cortical magnification function and that predicted from frequency discrimination. Future studies combining cortical and behavioural measurements within the same individuals will be important to confirm the present findings and to test whether individual differences in cortical magnification predict discrimination performance, as reported in vision and touch (Duncan and Boynton, 2007, 2003).

Second, our estimates of cortical magnification rely on BOLD fMRI, an indirect and spatially limited measure of neural activity. Although this is likely to broaden estimated tuning widths (Besle et al., 2022), it should have only a minor effect on preferred frequency estimates, and hence on the inferred frequency mapping. Consistent with this, previous work has shown good agreement between metabolic and electrophysiological estimates of best frequency (Kalatsky et al., 2005).

Third, some variability in the estimated magnification functions arose from methodological factors, including small variations in stimulus and scanning parameters across participant groups and the choice of analysis method. While the main conclusions were robust across these variations, this suggests some sensitivity of the exact magnification profile to measurement details. Observed differences between participant groups may reflect either methodological effects or sampling variability. Likewise, the three estimation approaches (gradient magnitude, cortical distance, and surface-based methods) yielded slightly different results, reflecting their distinct assumptions about tonotopic organisation. Notably, however, both linear methods (gradient magnitude and cortical distance) produced broadly similar estimates, suggesting that the overall magnitude of cortical magnification is relatively stable despite these differences.

A related issue concerns the precise functional status of the identified tonotopic gradients. All three magnification estimates assume that the underlying maps have been correctly localised, yet the parcellation of human auditory cortex remains debated (Moerel et al., 2014; Saenz and Langers, 2014). In the present data, we observed only minor differences in magnification and frequency mapping between the anterior and posterior gradients. This relative similarity is consistent with the possibility that both gradients belong to primary auditory cortex, in line with previous interpretations (Besle et al., 2019; Da Costa et al., 2011), or, alternatively, that cortical magnification does not vary systematically across hierarchical stages of auditory cortex. In non-primate models, magnification has been reported to be similar across distinct auditory fields (Storace et al., 2011), suggesting a degree of invariance at least at early cortical levels. However, other sensory systems highlight that seemingly stable magnification functions can coexist with substantial transformations of the underlying representations. In vision, for example, cortical magnification varies relatively little across early areas, yet receptive fields increase substantially in size, suggesting systematic transformations of the underlying representation (Harvey and Dumoulin, 2011). Characterising whether analogous representational changes also occur in auditory cortex will be important for understanding how transformations across cortical stages relate to perceptual performance.

## Conclusion

Overall, the fine-grained tonotopic organisation of primary auditory cortex was better predicted by frequency discrimination performance than by cochlear tonotopic organisation, suggesting that frequency discrimination may be constrained primarily by cortical rather than cochlear processes. This relationship between cortical magnification and hyperacuity performance parallels observations in the visual and tactile systems, where primary cortices selectively magnify behaviourally relevant regions of sensory space beyond peripheral representations, potentially supporting hyperacute perceptual judgements. Despite these parallels, important differences remain. In contrast to vision: auditory cortical magnification emphasises a range of frequencies that is not already over-represented in the periphery. This dissociation suggests that cortical magnification in audition is not simply inherited from peripheral encoding, but instead reflects central processing demands, highlighting a key role for cortical organisation in shaping sensory acuity.

## Methods

### Frequency mapping and magnification predictions

We predicted cortical frequency magnification and mapping functions from published psychophysical measurements of auditory filter widths and frequency discrimination thresholds as a function of frequency. For auditory filter widths, we used the function obtained by Oxenham and Shera (2003) based on notched-noise forward-masking measurements expressed as a power function of the signal frequency:

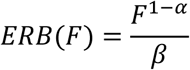

where ERB is the equivalent rectangular filter bandwidth (in kHz), F is the signal frequency in kHz, *α* = 0.27 and *β* = 11.1.

For frequency discrimination thresholds, we used the function obtained by Micheyl et al. (2012), who fitted log-transformed difference limens for frequency (DLFs) from 12 published pure-tone frequency discrimination studies with a linear combination of power functions of the reference tone’s frequency, duration and level:

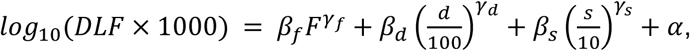

where *DLF* is the frequency discrimination threshold in kHz, *F* is the reference tone’s frequency in kHz, *d* is the tone duration in ms and *s* is the tone’s sensation level (in dB). The best-fitting parameter estimates were *γ_f_* = 0.82, *β_f_*= 0.38, *γ_d_* = -0.42, *β_d_* = 0.42, *γ_s_* = 0.82, *β_s_* = 0.37 and *α* = -0.38. For the current analysis, we set *d* = 200 ms and *s* = 40 dB SL (approximately corresponding to the stimulus duration and level used in our fMRI experiment), which simplifies the relationship to:

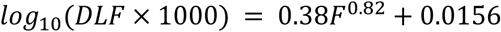

or:

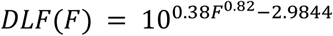

Under the assumption that psychophysical measures of frequency resolution or discrimination (ERBs or DLFs) correspond to a fixed cortical distance at any frequency, the predicted cortical frequency magnification function should be proportional to the reciprocal of the ERB or DLF function, referred to as ERB/DLF-rate function, and the predicted cortical frequency mapping function should be proportional to the antiderivative (integral) of the rate function, referred to as ERB/DLF-number function (Greenwood, 1990; Moore, 1986). For the ERB-based model, we derived the ERB-number function analytically:

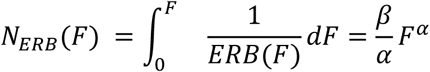

and for the DLF-based model, we estimated the DLF-number function, 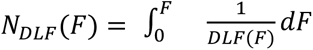, numerically using Matlab’s *integral* function.

In Figure 1, the predicted cortical magnification and mapping functions are expressed in mm/Hz and mm, respectively, and were obtained by multiplying the ERB/DLF rate and number functions by a scaling constant in ERB/mm or DLF/mm. The scaling constants were chosen to approximate our empirical fMRI-derived estimates of the cortical magnification and mapping functions (Fig. 4) and were set to 0.155 mm/ERB and 0.008 mm/DLF, respectively. In addition, the predicted mapping functions in Fig. 1C are plotted with an added constant, such that they have a value of 0 at 40 Hz instead of 0Hz (the value of the constant is immaterial, since the mapping function was later linearly fitted to the respective fMRI-derived functions with a free intercept parameter).

### Participants

This study involved 20 normal-hearing participants with no history of neurological or otological disease (8 males; mean age: 29.7 ± 7.4 years, SD). All participants gave informed written consent. The procedures complied with the Declaration of Helsinki guidelines (Version 6, 2008) and were approved by the Ethics Committee of the University of Nottingham’s School of Medicine. Data from participants 1 to 12 have been presented previously in a different form (Besle et al., 2022, 2019),

### Auditory stimuli

Auditory stimuli were generated in Matlab, digital-to-analog converted and time-controlled using a TDT System 3 with RP2.1 real-time processor (Tucker Davis Technology, Alachua, FL, USA), and presented binaurally using MRI-compatible insert earphones (participants 1-12: EarPlug, Nordic Neurolab, Bergen, Norway; participants 13-20: S14, Sensimetrics, Gloucester, MA, USA).

Stimuli were trains of narrowband noises (NBNs) at fixed centre frequencies, presented during the 6.25-s silent period between BOLD acquisitions (sparse sequence, see next section). For participants 1-12, NBNs were presented at one of seven centre frequencies between 0.25 and 6 kHz, with 24 trials per centre frequency (Fig. 2A). For participants 13-20, NBNs were presented at one of 32 centre frequencies between 0.1 and 8 kHz, with four trials per centre frequency. In both cases, NBN centre frequencies were equally spaced on an ERB-number scale derived from simultaneous notched-noise masking measurements (Glasberg and Moore, 1990). All NBNs had bandwidths of 1 ERB on that scale.

Each fMRI trial consisted of a ∼5-s train of NBN bursts starting after the end of each BOLD volume acquisition. Participants 1-6 were presented with trains of ten 200-ms NBN bursts separated by 300 ms intervals (total train duration: 4700 ms). For participants 7-12, each 200-ms burst was replaced by a 290-ms sequence of four 50-ms bursts, each separated by 30 ms (total trial duration: 4790 ms). Participants 13-20 were presented with a random sequence of sixteen 200-ms and ten 50-ms bursts, each separated by 50 ms intervals (total trial duration: 4950 ms). All NBN bursts were gated on and off with 10 ms quarter-cosine ramps.

NBNs at different frequencies were equalised to compensate for the frequency transfer characteristics of the earphones. All NBNs were presented at a root-mean-square level of 70 or 80 dB SPL (participants 1-12 and 13-20 respectively), above a continuous background of equally-exciting noise (Glasberg and Moore, 2000) set to 35 dB SPL per ERB, creating a constant hearing threshold of approximately 29 dB SPL at all frequencies (Hawkins and Stevens, 1950). Stimuli were therefore presented at approximately 41 and 51 dB SL for participants 1-12 and 13-20, respectively.

### Image acquisition

MRI data were collected on a Philips Achieva 7T scanner using a Nova Medical (Wilmington MA, USA) single-channel transmit, 32 channel receive (1Tx32Rx) head coil. Participants watched a self-chosen silent subtitled movie to stay alert and were instructed to ignore all sound stimuli.

BOLD fMRI data were acquired using a sparse (TR = 7.5 s) gradient-echo EPI sequence with 1.5 mm isotropic resolution (TE = 25 ms, SENSE = 3, FA = 90°, phase encoding direction: AP; Participants 1-12: 20 axial slices, FOV: 174 × 158 × 30 mm^3^ [AP × RL × FH], acquisition time = 1220 ms; participants 13-20: 24 axial slices, FOV: 156 × 174 × 36 mm^3^, acquisition time = 1250 ms). The acquisition stack was oriented parallel to the Sylvian fissure. The sound stimuli (NBN trains) started 1.28 s (participants 1-12) or 0.75 s (participants 13-20) after the end of the preceding acquisition and finished before the start of the next acquisition. For participants 1-12, data were acquired in six runs of eight minutes each, and each run contained four trials at each of the seven centre frequencies and 12 empty (baseline) trials. Each run also contained 24 additional trials with NBNs at variable centre frequencies within the same train, which were analysed as part of another publication (Besle et al., 2022). These additional trials were included in the General linear model (GLM) fit (see next section) but not considered further in the present report. For participants 13-20, data were acquired in two runs of nine minutes each, with each run consisting of two trials at each of the 32 centre frequencies and eight empty trials.

A whole-head, high-resolution T1-weighted structural volume was acquired using a phase-sensitive inversion recovery (PSIR) sequence (participants 1-12: TR = 15 ms, TE = 6.1 ms, FA = 8°, TI1/TI2 = 742/2685 ms, FOV = 200 × 181 × 140 mm^3^; participants 13-20: TR = 13 ms, TE = 3.76 ms, FA = 8°, TI1/TI2 = 791/2391 ms, FOV = 192 × 184 × 140 mm^3^) with 0.6, 1 or 0.8 mm isotropic resolution (participants 1-12, 13 and 14-20 respectively), which was used to create a 3D model of the supratemporal cortical surface in each participant (see Besle et al., 2019).

### Pre-processing and co-registration

Functional and structural data were pre-processed using a combination of FSL v6.0 (fsl.fmrib.ox.ac.uk/fsl/fslwiki/FSL; Jenkinson et al., 2012), MRTrix3 v3.0.4 (https://www.mrtrix.org; Tournier et al., 2019), Freesurfer v7.4 (surfer.nmr.mgh.harvard.edu; Dale et al., 1999), mrTools v4.7 (gru.stanford.edu/doku.php/mrtools/overview; Gardner et al., 2018) and in-house python and Matlab scripts.

The functional volumes were corrected for distortions due to dynamic B_0_ inhomogeneities (see Besle et al., 2019 for details), motion-corrected, high-pass filtered (cut-off 0.01 Hz), converted to percent signal change, and then concatenated across runs. BOLD frequency tuning curves were estimated voxelwise by fitting a GLM to the concatenated functional time series to obtain the BOLD response to each of the 7 or 32 frequency conditions (see Besle et al., 2019 for details).

The T1-weighted PSIR volume was segmented using Freesurfer to reconstruct individual three-dimensional inner and outer cortical surfaces. Surface meshes were interpolated at an additional 9 intermediate cortical depths, defining cortical depths between 0 (inner surface) and 100% (outer surface) in 10% steps. Freesurfer white matter and grey matter masks were visually inspected and manually corrected to exclude veins in the Sylvian fissure before surface reconstruction. Flattened cortical patches of the supratemporal plane were created using mrTools with a target 2D resolution of 0.33 mm at each cortical depth. To project the functional data onto the 3D surface meshes and flattened cortical patches, the distortion-corrected functional volumes were non-linearly registered with the T1-weighted PSIR volume (using an additional T2*-weighted FLASH volume with the same slice prescription as the functional volumes in an intermediate registration step) (see Besle et al., 2019 for details).

Further analysis of the functional data was restricted to a region encompassing auditory-responsive voxels within the cortical sheet of the supratemporal plane in each hemisphere of each participant. This was done by thresholding voxels responding significantly to a least one frequency condition compared to baseline (p < 0.05, FDR-corrected across all functional voxels, Benjamini et al., 2006), displaying the thresholded voxels on the flattened cortical patch (averaged between 20 and 80% cortical depths), and manually drawing a convex ROI around the largest cluster of significant voxels overlapping Heschl’s gyrus. This ROI was then expanded orthogonally to the cortical surface to include the full cortical depth and voxels within 30% of total cortical thickness below and above the cortical sheet, excluding non-adjoining cortical regions).

### Tonotopic gradient ROI definition

Preferred frequencies and population frequency tuning widths were estimated at each voxel of the flattened cortical patch by fitting the GLM-derived vertex-wise BOLD frequency tuning curve with a Gaussian function of stimulus frequency expressed in the original stimulus frequency scale (the ERB-number scale). Preferred frequencies and frequency tuning widths were defined as the mode and FWHM of the best fitting Gaussians, respectively. Before fitting, the data were first smoothed along the cortical surface at each cortical depth with a 1.33-mm-FWHM 2D Gaussian kernel.

Reversals in the preferred frequency gradients were identified on the 2D flattened cortical space using an unbiased automated procedure (Besle et al., 2019; Schönwiesner et al., 2015). For this, the preferred frequency data were averaged across all cortical depths and further smoothed along the cortical surface using a 6-mm-FWHM Gaussian 2D kernel. In all 40 hemispheres, the automated procedure identified a low-frequency gradient reversal approximately aligned with Heschl’s gyrus, flanked anteriorly and posteriorly by two approximately parallel high-frequency reversals. In cases where the automated procedure detected separate but aligned reversals of the same polarity (low or high frequency), these were manually joined. The anterior and posterior gradient ROIs were defined as the areas of cortical surface between the low-frequency reversal and either of the two high-frequency reversals. Gradient ROIs were projected back from the flattened cortical patch to the 3D space of each individual brain for further analysis.

### Derivation of cortical frequency magnification and mapping functions

The three measurements used to characterise frequency mapping in the gradient ROIs (local frequency gradient magnitude, cortical surface dedicated to different preferred frequencies and cortical position along the tonotopic gradient) were estimated on the cortical surface mesh of individual participants. For all three methods, preferred frequencies were first re-computed at each vertex of the cortical surface mesh (instead of each voxel of the flattened cortical patch previously) by fitting Gaussians to the unsmoothed BOLD tuning curve averaged across the central 60% of the cortical sheet. Gaussians were fitted to the tuning curves after converting stimulus frequency to either a logarithmic frequency scale (octave scale) or a DLF number scale, depending on whether magnification was expressed in mm/Hz (Figs 3 and 4A-C) or mm/DLF (Figs 4D-F).

All three measures were binned according to preferred frequency. We used 24 contiguous preferred-frequency bins equally-spaced on a log-frequency scale from 0.1 to 8 kHz. For the first 12 participants, who only heard stimuli between 0.25 and 6 kHz, bins outside this range were excluded, resulting in 17 bins. Binning allowed us to easily obtain frequency mapping or magnification functions by numerical differentiation/integration, to average frequency mapping functions across participants and to conduct statistical analyses across participants (see later sections).

#### Gradient magnitude

We derived the magnitude of the local preferred-frequency gradient (in Hz/mm or DLF/mm) at the centre of each cortical mesh triangle from the coordinates and preferred frequencies of its three vertices (Mancinelli et al., 2019). Gradient magnitudes were then binned by preferred frequency, estimated for each triangle by linear interpolation of the three vertex preferred frequencies. Frequency magnification was calculated as the reciprocal of the median gradient magnitude within each bin (in mm/Hz or mm/DLF). For a similar approach in the primary visual cortex, see Harvey and Dumoulin (2011).

#### Cortical surface

Frequency magnification at each preferred frequency bin was calculated as the total surface area (in mm^2^) of all cortical mesh triangles within a given gradient ROI whose preferred frequency fell within that bin, divided by the bin width (in Hz or DLFs).

#### Cortical distance

Cortical position along the tonotopic gradient was estimated as the geodesic distance between a given vertex within a gradient ROI and the low-frequency reversal marking the boundary between the two gradient ROIs. This was calculated as the shortest distance along the mid-cortical mesh from that vertex to any vertex within the low-frequency reversal. This method therefore assumes that the tonotopic gradient follows the shortest path between the low-frequency and high-frequency reversals in each gradient ROI. To account for variability in reversal distance both within gradient ROIs (low- and high-frequency reversals were often closer medially than laterally) and across individuals, we expressed the cortical distance of each vertex as a proportion of the shortest cortical distance between the low- and high-frequency reversals going through that vertex. The frequency mapping function was calculated by first binning the normalised cortical distances according to the preferred frequency of each vertex, taking the median normalised cortical distance in each bin, and multiplying it by the group-average reversal distance in the ROI to obtain cortical distances in mm as a function of preferred frequency. The average reversal distance for a given gradient ROI was computed as the average of all the shortest paths between vertices of the low- and high-frequency reversals. The group-average reversal distances were 11.5 ± 0.392 and 10.9 ± 0.446 mm for the left anterior and posterior gradients respectively, and 11.4 ± 0.344 and 10.9 ± 0.348 mm for the right anterior and posterior gradients. A Bayesian ANOVA provided anecdotal-to-moderate evidence against an effect of group (participants 1-12 vs 13-20), gradient (anterior vs posterior), hemisphere, or their interactions on reversal distance (all BF_10_‘s < 0.856). Distance-derived cortical magnification functions were computed by numerically differentiating the binned cortical distance with respect to bin centre frequency. For a similar approach in the primary visual cortex, see Engel et al. (1997). To end up with 24 bins after differentiation, cortical distances were binned using 25 bin centres between 0.1 and 8 kHz. To ensure positive cortical magnification values, the cortical frequency mapping function was first fitted with a monotonically increasing spline function (using the Shape Language Modelling toolbox for Matlab).

### Statistical analyses

For all analyses involving frequency-binned data points, we excluded outliers (participants) at each preferred frequency bin if the measurement was more than 3×1.4826 median absolute deviations (equivalent to 3 standard deviations) away from the median measurement for this bin. This was done separately for each preferred frequency bin, gradient ROI and hemisphere.

Linear transformations of the ERB- and DLF-predicted functions were fitted to the empirical fMRI-derived functions by minimizing ordinary least squares. For gradient magnitude functions expressed in Hz/mm and magnification functions expressed in mm^(2)^/Hz, the empirical data and predictions were log-transformed before fitting. Best linear fits for the ERB and DLF predictions were compared using the Bayes information criterion (BIC) approximation of the Bayes factor (BF; Raftery, 1995; Wagenmakers, 2007), with BF_DLF,ERB_ larger than 1 providing evidence for the DLF scale and BF_DLF,ERB_ smaller than 1 providing evidence for the ERB scale. The strength of evidence provided by BFs was interpreted following Kass and Raftery (1995): BF between ⅓ and 3: anecdotal/weak evidence; BF below ⅓ or above 3: moderate/positive evidence; BF below 0.05 or above 20: strong evidence; BF below 0.0067 or above 150: very strong evidence. The BIC was computed from the residuals of a single linear fit to all individual empirical functions within a group of participants. The number of observations for computing the BIC (12 × 17 = 204 and 8 × 24 = 192 data points for participants 1-12 and 13-20 respectively) was adjusted to account for correlations between preferred frequency bins. This was done by replacing the number of data points by the effective sample size (Thiébaux and Zwiers, 1984), computed by dividing the number of data points by the sum of the autocorrelation function of the residuals across preferred frequency bins. For a given comparison, we used the average effective sample size across the ERB and DLF linear fits. Average effective sample sizes ranged between 36.7 and 44.4 (participants 1-12) and between 27.5 and 42.6 (participants 13-20) across gradient magnitude, cortical surface and cortical distance functions.

All inferential hypothesis tests were conducted using the BayesFactor package in R (v4.2.3), with the default Jeffreys-Zellner-Siow (JZS) prior (a Cauchy distribution with scale parameter 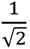) for mixed-effect ANOVA and *t*-tests (Rouder et al., 2012, 2009). The strength of evidence provided by BFs were interpreted following Kass and Raftery (1995; see previous paragraph), with the convention that BF_10_ above 1 provides evidence for the alternative hypothesis (H_1_) against the null (H_0_). Effects were only reported, and interaction effects decomposed into simple effects, if the corresponding BF_10_ provided at least moderate evidence for H_1_ (BF_10_ > 3), unless stated otherwise.

BFs were adjusted from multiple comparisons according to Jeffreys’ method (Jeffreys, 1938) when testing multiple independent hypotheses (e.g. testing an effect at each of the 24 preferred frequency bins). Briefly, instead of assuming that both H_0_ and H_1_ have the same prior probability (0.5) for each comparison when interpreting the evidence given by the BF, the prior probability of each comparison’s H_0_ was increased such that the prior probability of all comparisons’ H_0_s being simultaneously true is equal to the prior probability of at least one H_0_ being false (both equal to 0.5). Each comparison’s prior probability for H_0_ was therefore increased from 0.5 to p = (0.5)^1/k^, where *k* = 24 is the number of independent comparisons (de Jong, 2019; van den Bergh et al., 2020). We then calculated the posterior odds of each comparison by multiplying its uncorrected BF_10_ by its prior odds p(H_1_)/p(H_0_) = (1-p)/p. Adjusted prior odds for the null hypothesis were 34.13 for independent tests across 24 bins. Posterior odds for a given comparison can be conceived as corrected BFs across all tested comparisons (Williams et al., 2016). This method is akin to Bonferroni correction in classical null hypothesis significance testing (de Jong, 2019). Since Bonferroni correction is overly conservative when the tests are correlated, we report both the uncorrected and corrected BFs.

All reported error intervals represent the parametric standard error of the mean, unless stated otherwise.

## Data availability statement

The data that support the findings of this study are openly available in openNeuro at https://openneuro.org/datasets/ds008460. Analysis code used to generate the results presented in this study is available at GitHub (https://github.com/julienbesle/frequencyCorticalMagnification7T) and archived at Zenodo (DOI: https://doi.org/10.5281/zenodo.21879249, version 0.1.1).

## Acknowledgement

This work was supported by the UK Medical Research Council [grant numbers G0901321, MC_U135097128, MR/S003320/1, MC_UU_00010/2] and by the University Research Board of the American University of Beirut.

## Figures supplements

### Figure 2 supplements

Figure 2 supplements 1 and 2 show the preferred frequency maps, tuning width maps and gradient ROIs for all participants.

**Figure S2.1:**
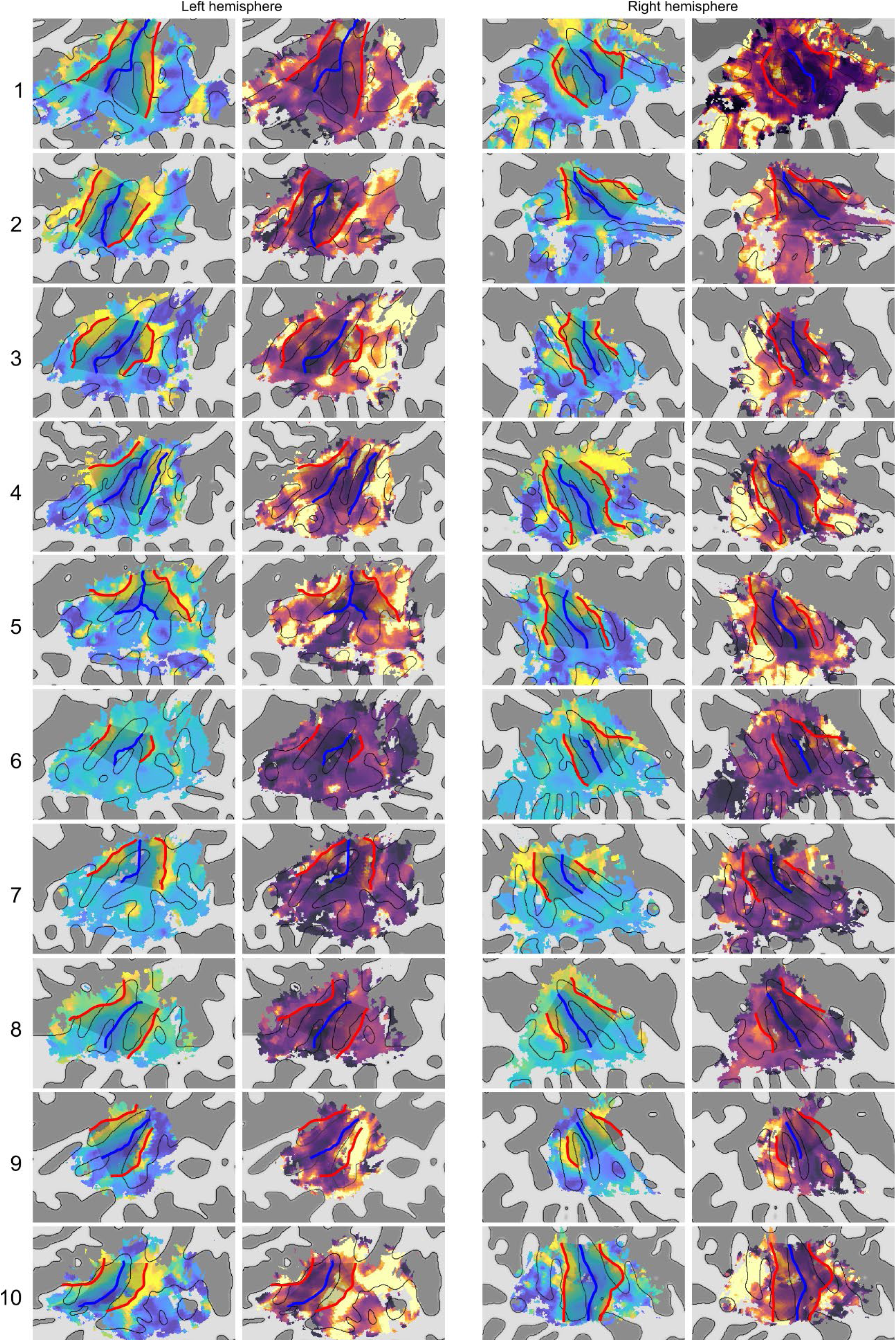
Preferred frequency and frequency tuning width in each hemisphere of participants 1 to 10.

**Figure S2.2:**
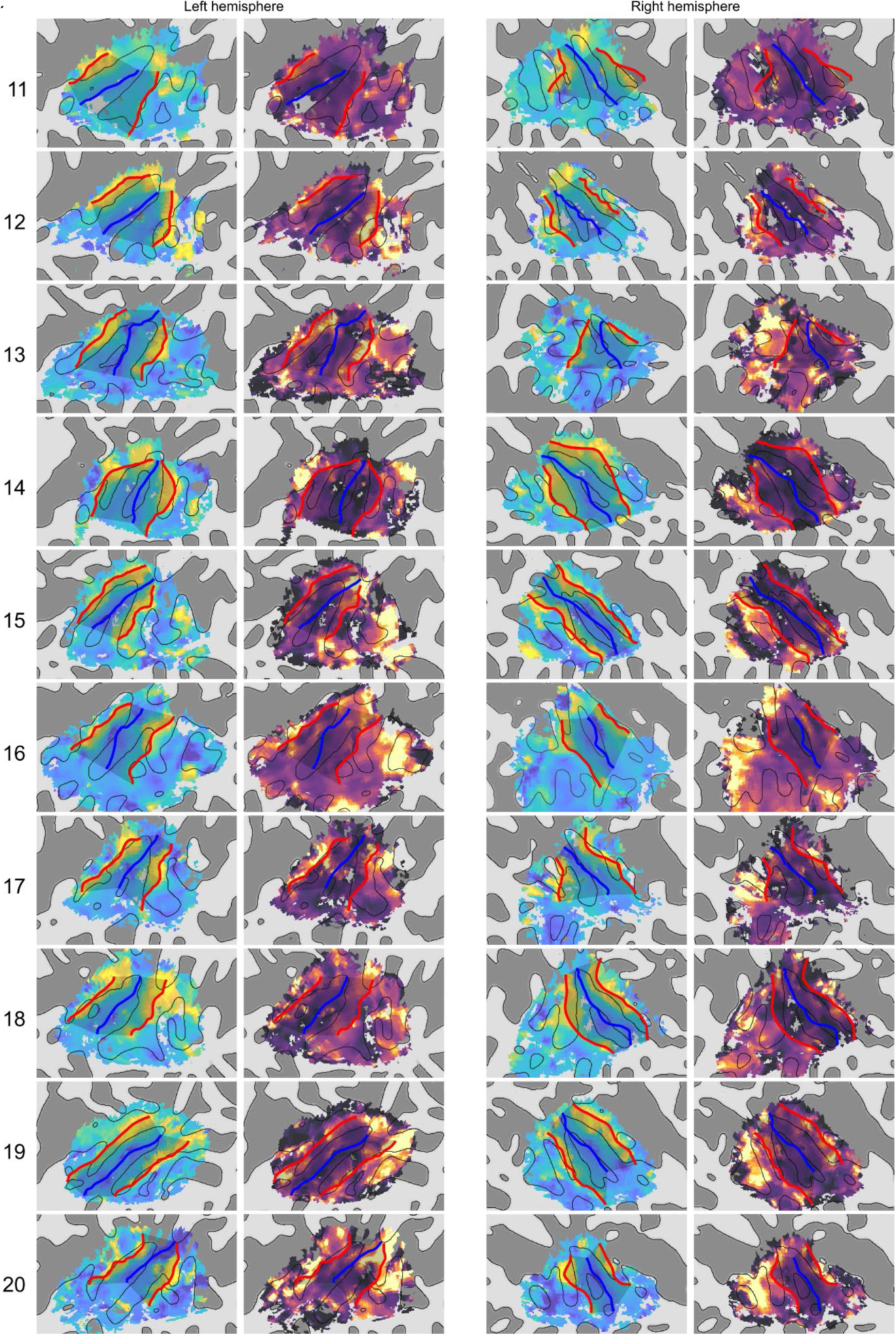
Preferred frequency and frequency tuning width in each hemisphere of participants 11 to 20.

### Figure 3 supplements: ERB and DLF model fit comparisons in separate gradient ROIs and hemispheres

In the main results, we fit the ERB- and DLF-based predictions to gradient magnitude, cortical surface and cortical distance data averaged across both hemispheres and both gradient ROI (separately for each group of participants) and find that the DLF model fits the data better than the ERB model for all three measurements and both participant groups. Here we show the same three measurements averaged separately in each hemisphere (Fig. S3A-C) and separately in each gradient ROI (Fig. S3D-F) (and averaged across all 20 participants), as well as the corresponding best-fitting ERB- and DLF-based predictions.

The resulting gradient, magnification and mapping functions were very similar both across the left and right hemispheres and across the anterior and posterior gradient ROIs (see next section for a statistical comparison showing minor differences). Bayesian model comparisons using the Bayesian information criterion approximation provided very strong evidence in favour of the DLF model (compared to the ERB model) in both hemispheres and in both gradient ROIS, for all three measures (*gradient magnitude*: left hemisphere: BF_DLF,ERB_ = 4.43 × 10^13^, right hemisphere: BF_DLF,ERB_ = 3.11 × 10^12^, anterior gradient: BF_DLF,ERB_ = 9.70 × 10^12^, posterior gradient: BF_DLF,ERB_ = 1.27 × 10^14^; *cortical surface*: left hemisphere: BF_DLF,ERB_ = 5.25 × 10^6^, right hemisphere: BF_DLF,ERB_ = 1.35 × 10^8^, anterior gradient: BF_DLF,ERB_ = 2.03 × 10^7^, posterior gradient: BF_DLF,ERB_ = 1.58 × 10^11^ ; *cortical distance*: left hemisphere: BF_DLF,ERB_ = 1.08 × 10^5^, right hemisphere: BF_DLF,ERB_ = 1177, anterior gradient: BF_DLF,ERB_ = 6180, posterior gradient: BF_DLF,ERB_ = 1.04 × 10^5^).

**Figure S3.1:**
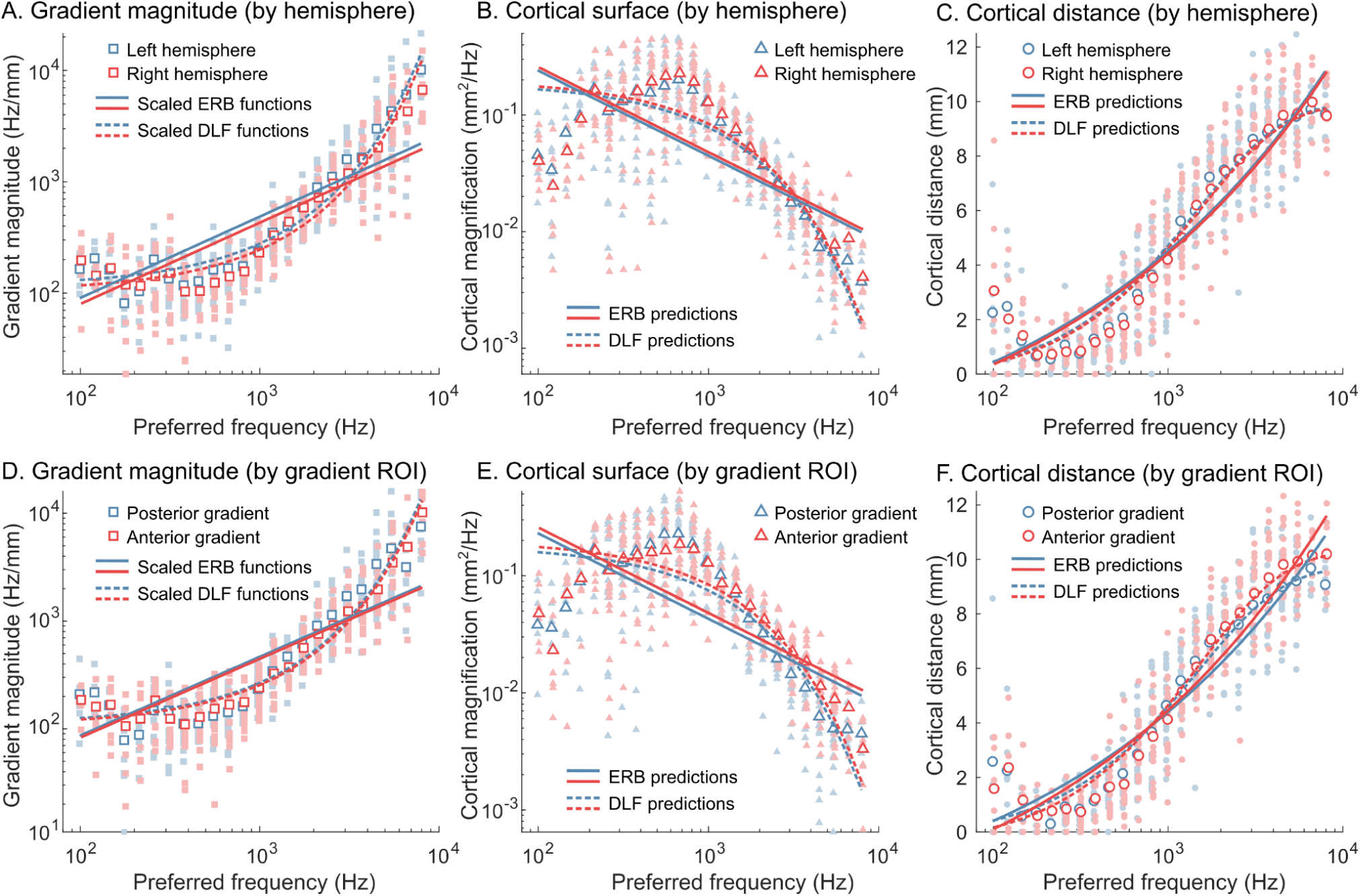
Gradient magnitude (A,D), cortical surface (B,E) and cortical distance (C,F) plotted as a function of preferred frequency, averaged across all 20 participants, separately for the left and right hemispheres (A-C) or for the anterior and posterior gradients (D-F). Plotting conventions are otherwise identical to those of Figure 3.

### Figure 4 supplements: Differences in cortical magnification estimates between gradient ROIs and hemispheres

In the main results, we report the results of a Bayesian ANOVA testing the effect of preferred frequency, group, hemisphere and gradient ROI on cortical frequency magnification measurements (expressed as cortical distance per DLF) for each of the three measurement methods (gradient magnitude, cortical surface or cortical distance). Effects of preferred frequency and group were fairly consistent between methods and are reported in the main results section. Here we report additional evidence for differences in frequency magnification between gradient ROIs and hemispheres and any interaction of these variables with group or preferred frequency. These effects were either small or restricted to a few frequency bins, and were inconsistent between measurement methods.

#### Effect of hemisphere

Both the gradient magnitude and cortical surface measurements provided at least moderate evidence for a main effect of hemisphere, with cortical magnification slightly larger on average in the right than in the left hemisphere (Fig. S4A-B; gradient magnitude: BF_10_ = 452, 10.3 ± 0.245 vs 8.90 ± 0.245 μm/DLF; cortical surface: BF_10_ = 7.66, 0.223 ± 0.00721 vs 0.195 ± 0.00648 mm^2^/DLF). However the cortical distance measurements provided moderate evidence against an effect of hemisphere (Fig. S4C; BF_10_ = 0.0622). There was no evidence for any interaction effect involving factor hemisphere.

**Figure S4.1:**
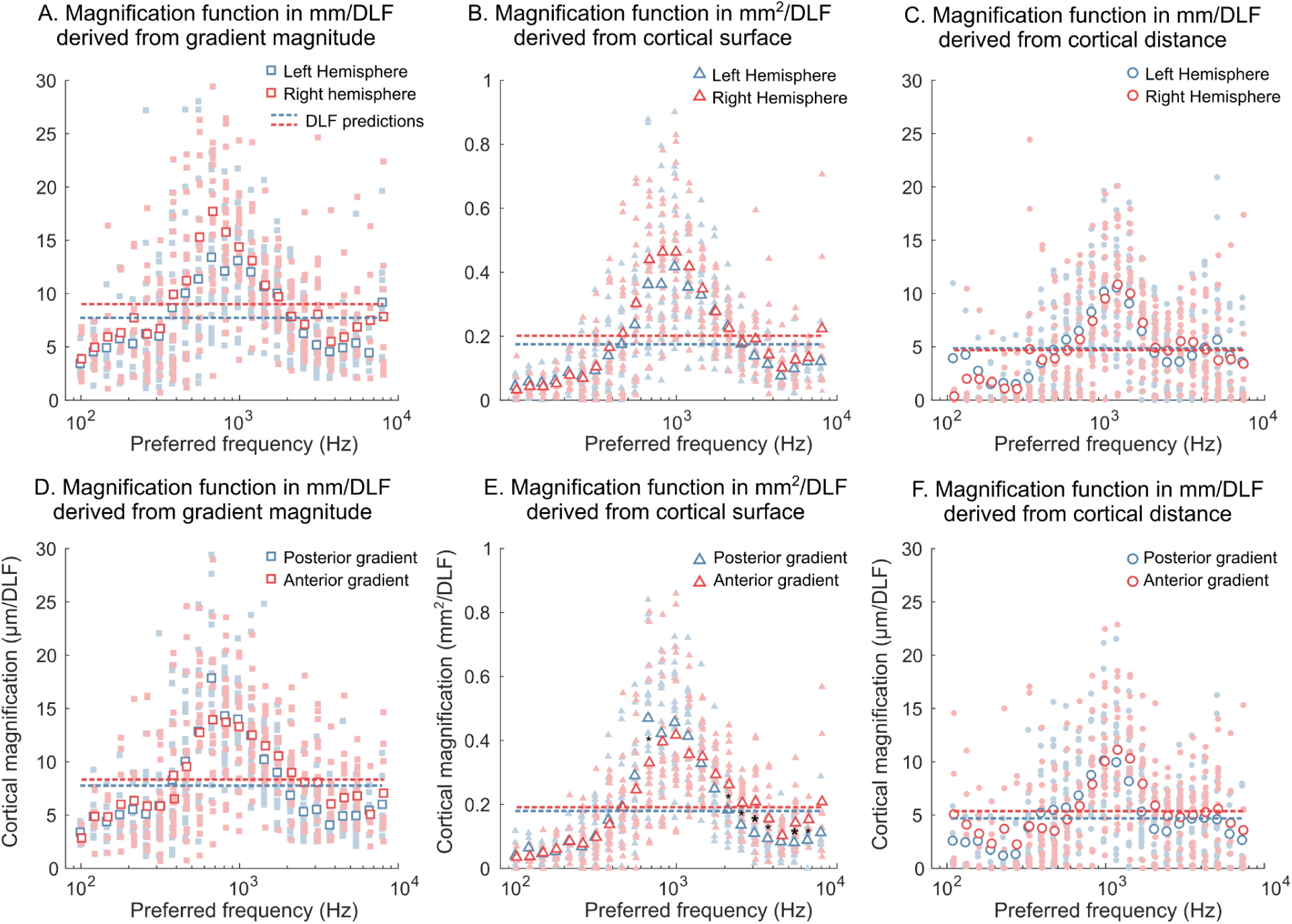
Differences in frequency magnification between left and right hemispheres (A-C) and between anterior and posterior gradient ROIs (D-F). A. Frequency magnification in mm/DLF, computed as the reciprocal of the local preferred frequency gradient magnitude, plotted as a function of preferred frequency and averaged separately for the left and right hemispheres (blue and red symbols, respectively). Open symbols show group averages and filled symbols show individual data. The dashed lines depict the best-fitting DLF predictions for each hemisphere, which also correspond to the average cortical magnification across preferred frequencies. B. Same as A, but frequency magnification was estimated as the surface area of cortical mesh vertices preferring different frequencies. C. Same as A, but frequency magnification was estimated by differentiating cortical distance along the tonotopic gradient with respect to preferred frequency. D-F. Same as A-C, but cortical magnification was averaged separately for the posterior and anterior gradient ROIs (blue and red symbols, respectively). In E, small and large black star symbols indicate preferred frequency bins where there was at least moderate evidence (uncorrected and corrected respectively) that cortical magnification differed between the two gradient ROIs.

#### Effect of gradient ROI

All three measurements provided moderate evidence against a main effect of gradient ROI, suggesting that cortical magnification did not differ between the anterior and posterior gradient ROIs, when averaged across frequencies/cortical locations (Fig. S4D-F; BF_10_ = 0.265, 0.112 and 0.141, respectively). However, both the gradient magnitude and the cortical surface measurements provided evidence for an interaction between gradient ROI and other factors. For gradient magnitude, there was moderate evidence for a three-way interaction between gradient, preferred frequency and group (BF_10_ = 5.41). This interaction was due to an interaction between gradient and preferred frequency in participants 1-12 (BF_10_ = 6.12 × 10^6^), but not participants 13-20 (BF_10_ = 0.00188). For participants 1-12, post-hoc T-tests at each frequency bin provided at least moderate evidence that frequency magnification was larger in the posterior than in the anterior gradient ROI at one low-frequency bin (0.61-0.71 kHz; uncorrected BF_10_ = 39.6) and larger in the anterior than in the posterior gradient ROI at six high frequency bins between 1.92 and 6.01 kHz (all uncorrected BFs > 4.01). None of these differences survived correction for multiple tests across frequencies. For cortical surface measurements, there was very strong evidence for an interaction between gradient ROI and preferred frequency (Fig. S3E; BF_10_ = 1.98 × 10^3^). Post-hoc tests provided evidence that frequency magnification was larger in the posterior than in the anterior gradient at one low-frequency bin (0.61-0.71 kHz; uncorrected BF_10_ = 21.1) and larger in the anterior than in the posterior gradient at six high frequency bins between 1.92 and 7.27 kHz (all uncorrected BFs > 8.53). The latter effect survived correction for multiple tests at 2 frequency bins (both corrected BFs > 9.49). Finally, the cortical distance measurements provided strong evidence against both an interaction between gradient ROI and preferred frequency (BF_10_ = 0.00461) and a three-way interaction between gradient ROI, preferred frequency and group (BF_10_ = 0.0124). There was no evidence for any other interaction effect involving factor gradient ROI.

